# Maternal- and Somatic-type snoRNA Expression and Processing in Zebrafish Development

**DOI:** 10.1101/858936

**Authors:** Johanna F. B. Pagano, Mauro D. Locati, Wim A. Ensink, Marina van Olst, Selina van Leeuwen, Wim C. de Leeuw, Ulrike Nehrdich, Herman P. Spaink, Han Rauwerda, Martijs J. Jonker, Rob J. Dekker, Timo M. Breit

## Abstract

Small nucleolar RNAs (snoRNAs) are non-coding RNAs that play an important role in the complex maturation process of ribosomal RNAs (rRNAs). SnoRNAs are categorized in classes, with each class member having several variants present in a genome. Similar to our finding of specific rRNA expression types in zebrafish embryogenesis, we discovered preferential maternal- and somatic-expression for snoRNAs. Most snoRNAs and their variants have higher expression levels in somatic tissues than in eggs, yet we identified three snoRNAs; U3, U8 and snoZ30 of which specific variants show maternal- or somatic-type expression. For U3 and U8 we also found small-derived snoRNAs that lack their 5’ rRNA recognition part and are essentially Domain II hairpin structures (U-DII). These U-DII snoRNAs from variants showed similar preferential expression, in which maternal-type variants are prominently expressed in eggs and subsequently replaced by a somatic-type variants during embryogenesis. This differential expression is related to the organization in tandem repeats (maternal type) or solitary (somatic-type) genes of the involved U snoRNA loci. The collective data showed convincingly that the preferential expression of snoRNAs is achieved by transcription regulation, as well as through RNA processing. Finally, we observed small-RNAs derived from internal transcribed spacers (ITSs) of a U3 snoRNA loci that via complementarity binding, may be involved in the biosynthesis of U3-DII snoRNAs. Altogether, the here described maternal- and somatic-type snoRNAs are the latest addition to the developing story about the dual ribosome system in zebrafish development.

## INTRODUCTION

Small nucleolar RNAs (snoRNAs) are a class of non-coding RNA molecules of variable length (the majority being 60-200 nucleotides long), found in archaea and eukaryotes (1). SnoRNAs are thought to mainly be involved in post-transcriptional modifications and maturation of ribosomal RNAs (rRNAs) (2,3). However, recently additional functions have been ascribed to specific snoRNAs, from regulation of mRNA editing and splicing (4) to post-transcriptional gene silencing (5,6). SnoRNAs do not possess any intrinsic catalytic or modification activity, but act both as a scaffold for partner proteins, forming small nucleolar ribonucleoproteins (snoRNPs) and as guide for target specificity (7). Based on base-pairing interactions with their target RNA, snoRNAs can thus direct the associated catalytic protein subunits to accurately modify a specific RNA site (8).

In general, eukaryotic genomes can contain up to 200+ unique snoRNA genes (9). Based on the presence of conserved sequence motifs, the majority of snoRNAs are classified into two distinct classes: box C/D snoRNAs, which guide 2’-O-methylation of ribose, and box H/ACA snoRNAs, which are involved in the isomerization of specific uridine residues to pseudouridine (1). C/D snoRNAs are defined by the presence of two conserved motifs, the C box (UGAUGA) and the D box (CUGA), found near the 5’- and 3’-end, respectively (10). In the folded C/D snoRNA molecule, these two motifs are in close proximity of each other by means of a hairpin structure and serve as a binding site for interacting proteins (11–13). In addition, many C/D box snoRNAs may have less-well-conserved copies of the C and D motifs (called C' and D'), which are also involved in the interaction with specific proteins (14–16). A conserved region of 7-20 nucleotides upstream of the D (and the D’, if present) box interacts via base-complementarity with the rRNA, pinpointing the to-be-methylated RNA base, which is usually the 5^th^ nucleotide from the D or D’ box (17–20). Additional base interactions with the target rRNA can stimulate methylation by up to five-fold (16). H/ACA snoRNAs typically feature a secondary structure consisting of two hairpins linked by a hinge region that contains the H-box (ANANNA), followed by a short tail with the ACA-box (ACA) (21). The hairpin regions contain internal bulges known as pseudouridylation pockets in which a conserved region of 6-20 nucleotides is complementary to the target, but leaves a uridine residue unpaired, marking it for enzymatic modification (22,23). However, more and more snoRNAs are discovered that don't follow this classification. For example, there are snoRNAs that have both C/D box and H/ACA box (24,25), or C/D snoRNAs that are not involved in 2’- O-methylation, like snoRNA U3 and U8, which instead are essential in the processing of 18S and 28S, 5.8S rRNAs, respectively (26–28).

Most vertebrate snoRNA genes are located in introns of genes that usually encode for proteins related to ribosome biogenesis and protein synthesis (29) (Figure 1A). After transcription of these genes by RNA polymerase II, the formation of functional snoRNAs requires the processing of intronic RNA sequences, which are released during pre-mRNA splicing (2). In vertebrates, only a few snoRNAs genes have an independent promoter and are directly transcribed by RNA polymerase II or III (30,31). snoRNA genes are present either as solitary snoRNA genes, or as clusters of multiple coding units; such clusters can consist of the same or different snoRNA genes (9) (Figure 1A).

**Figure 1.**
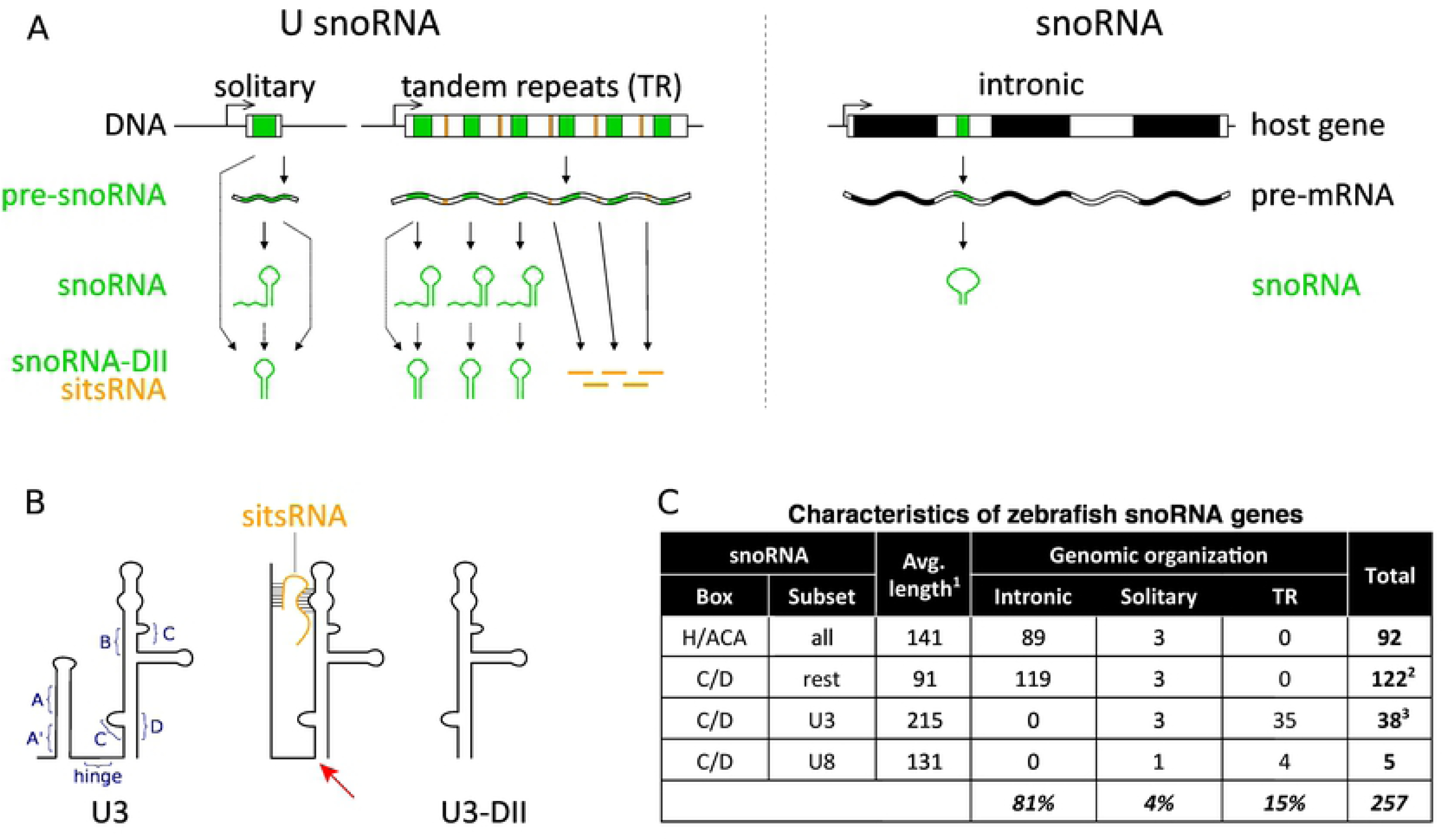
Genomic organization and RNA molecules from zebrafish snoRNA genes. **A:** Different configurations of genomics organization of snoRNA genes illustrated by exemplifying variants (upper row) and the different forms of RNA transcripts (middle rows). For the U snoRNA genes also derived transcripts are indicated (lower row): snoRNA-DII (green) are transcripts that exist just of the Domain-II part of the U snoRNA gene; sitsRNA (yellow) are snoRNA-ITS-derived-small-RNAs that originate from ITS regions in clusters of the tandemly repeated U snoRNA genes. The dotted lines indicate possible RNA processing routes for snoRNA-DII transcripts. **B:** The various snoRNA boxes are indicated in a schematic representation of the U3 snoRNA (left panel). The possible interaction of a chromosome 14 U3 sitsRNA with a complete U3 snoRNA is indicated (middle panel), which may play a role in processing the latter at the position of indicated by the red arrow to produce a U3-DII transcript. A schematic representation of the U3-DII transcript found in this study (right panel). **C:** Several characteristics of zebrafish snoRNA genes organized by their H/ACA or CD boxes. ^1^Distribution plot of lengths of snoRNA genes (Supplemental Figure SF1); ^2^snoU2_19 and SNORD94_a are present twice in the genome; ^3^U3_l and U3_k were found two and five times in the genome, respectively.

An increasing number of recently identified snoRNAs show differential expression among different cell types and tissues, suggesting a role in distinct physiological processes (6). For example, several studies discovered that certain snoRNAs are expressed mainly in the brain where they might affect organ function and/or development (32,33).

Our group has recently described the presence of two distinct types of rRNAs expressed during zebrafish development: a maternal-type, exclusively accumulated during oogenesis, and replaced throughout the embryogenesis by a somatic-type, which is the only one rRNA type present in adult somatic tissue (34,35). Since maternal- and somatic-type rRNAs show ample sequence differences combined with the fact that snoRNAs interact with rRNA in a sequence specific way, we hypothesized that there also might exist specific maternal- and somatic-type snoRNAs. For this, we investigated the expression of the snoRNAome throughout zebrafish development from egg to adult, by small-RNA-seq and discovered that indeed complete and partial transcripts of snoRNA variants exist that show either maternal- or somatic-preferential expression. Moreover, we determined that this developmentally regulated expression is likely regulated both at the level of transcription, as well as RNA processing. Together, with our recent finding that also a maternal-type spliceosome with specific snRNA variants exists for early embryogenesis (36) everything points to the existence of a comprehensive dual translation system in zebrafish embryogenesis.

## RESULTS AND DISCUSSION

### Cataloguing the zebrafish snoRNAs

In our previous studies (34,35), we identified two distinct types of rRNAs in zebrafish development: maternal and somatic. The maternal-type rRNAs make up virtually all the rRNA present in oocytes and are gradually, yet completely, replaced by the somatic-type rRNAs during embryogenesis. There are, for each rRNA species, 5S, 5.8S, 18S and 28S, significant sequence differences between the two types of rRNA, indicative of a substantial functional difference. Given that snoRNAs are intricately involved in the complex maturation process of rRNA via sequence specific interactions, we investigated whether there are specific snoRNAs being co-expressed with maternal- and somatic-type rRNAs.

As the zebrafish genome is quite well annotated, we started by making an inventory of the known genomic snoRNA sequences. In general, snoRNAs are categorized in two main families based on the presence of conserved sequence motifs: the C/D box and H/ACA box (37) and they can be found in the databases snOPY (38) and Ensemble 89 (39) (Supplemental Table ST1 and Supplemental Figure SF1). Collectively, 257 snoRNA loci were identified on the 25 zebrafish chromosomes. However, since some of these loci contained identical sequences, 250 zebrafish snoRNA sequences were found to be unique (Supplemental File SF1). In contrast, several snoRNAs appear to be present as different variants, which we labeled with a unique identifier (Supplemental Table ST1). Most snoRNA loci are present in introns (*intronic, 81%*) and rely for their expression on the transcription of the associated genes (Figure 1A and 1C). Yet, a few snoRNA loci, mainly belonging to the C/D box snoRNAs, are expressed as independent transcriptional units, consisting of one (*4%*) or several (*15%*) snoRNA genes (Figure 1A and 1C, Table 1 and Supplemental Figure SF1).

Most transcriptionally self-regulated snoRNAs have been named U3 and U8 (40). The snoRNA loci appear to be non-randomly distributed over the chromosomes, but no apparent co-location with known rRNA sequences was observed (Supplemental Table ST1).

### Developmental-stage-specific expression of two snoRNA types

After the identification and annotation of all snoRNAs in the zebrafish genome (GRCz10), we investigated whether, similar to rRNAs, maternal- and somatic-type snoRNAs exist. For this, we analyzed the differential expression between egg and adult zebrafish of the 204 expressed snoRNAs found by small-RNA-seq. The distribution of snoRNAs based on their differential expression was clearly bimodal, with no snoRNA-variant being equally expressed in egg and adult zebrafish (Figure 2A). Hence, all these snoRNAs displayed preferential expression, as either maternal- (n = 18) or somatic-type (n = 186) snoRNA (Figure 2), albeit less absolute as previously observed with maternal- and somatic-type rRNAs (Supplemental Table ST2). The maternal-type snoRNAs turned out to be variants of just three snoRNAs: U3, U8 and snoZ30, each of which also has one somatic-type variant (Figure 2B). Remarkably, for the U3 and U8 snoRNAs, the variants in tandem repeats showed a maternal-preferential expression profile, whereas the solitary variants were somatic-preferential (Figure 2B). For both U3 and U8 in tandem repeats, there were some variants that showed no expression (Supplemental Figure SF2). For the snoZ30 snoRNAs, the maternal- and somatic-type variants are also organized in a different way on the genome with the maternal-type being a solitary snoRNA, whereas the somatic-type is located intronically (Figure 2B). Thus, there is a clear association between the genome organization of the snoRNA variants and their expression during development.

**Figure 2.**
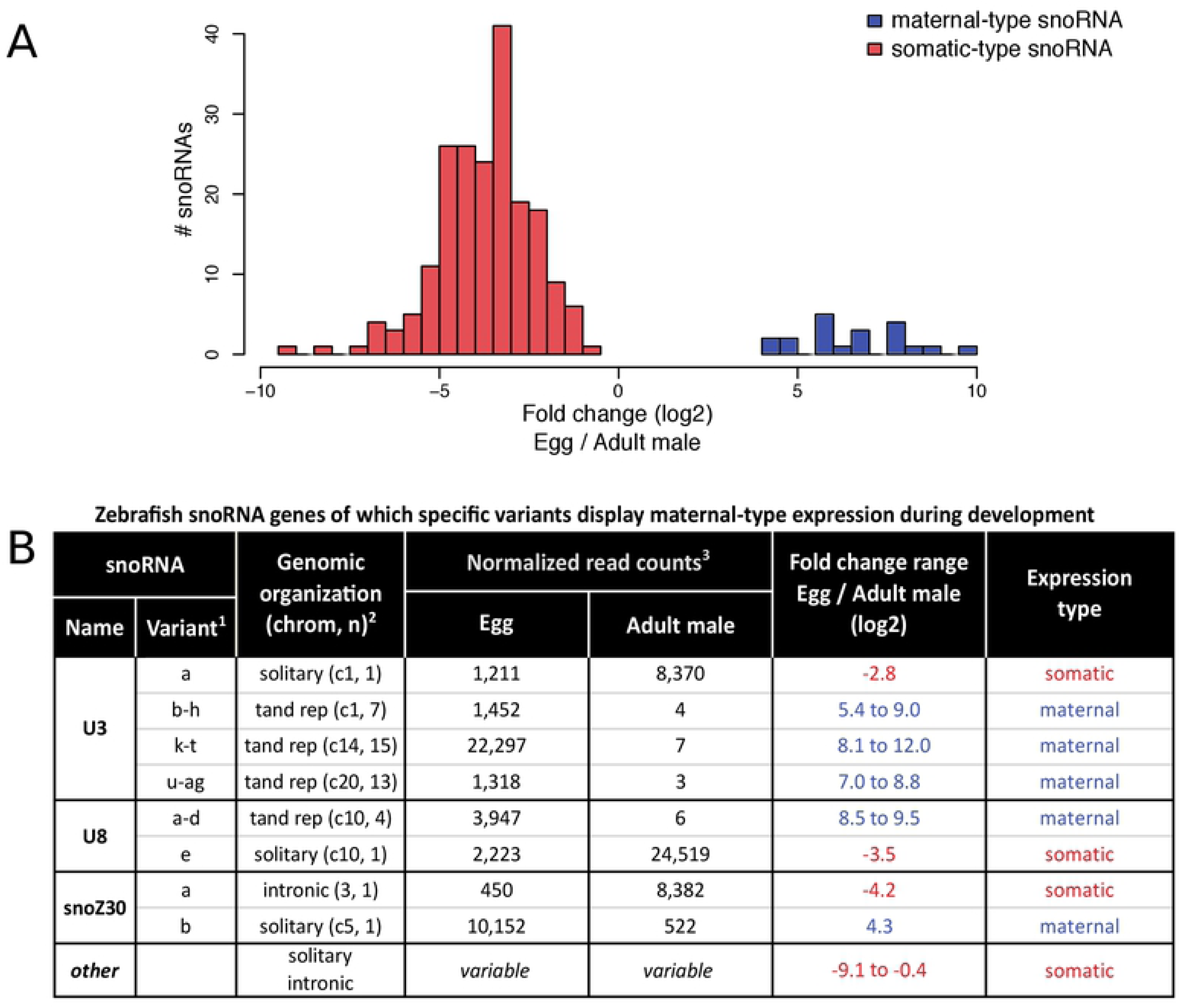
Identifying maternal and somatic types snoRNAs. **A:** Distribution plot of differential snoRNA expression between zebrafish egg and adult-male tissue. Maternal-type snoRNA (blue), 18 snoRNA variants with a positive fold change, i.e. with a higher expression in egg than in adult-male tissue; somatic-type snoRNA (red), 186 snoRNA variants with a negative fold change. **B:** Overview of maternal- and somatic-type snoRNA variants of U3, U8 and snoZ30 genes. ^1^Names of the associated variants within each of the indicated snoRNA locus. ^2^Genomic organization: intronic, solitary or in clusters of tandemly repeated (tand rep) genes. Chromosome number (chrom) plus the number (n) of variant genes in a snoRNA locus. ^3^Normalized NGS-read counts of the snoRNA variants (cf. Supplemental Table ST2) in the same cluster of tandemly repeated genes were added up.

There are ample sequence differences between the maternal and somatic snoRNA types (Figure 3A). We used these sequence differences to confirm the developmental-stage preferential expression of maternal- and somatic-type snoRNAs, by qRT-PCR-analysis for several selected maternal- and somatic-type U3, U8 and snoZ30 snoRNAs in egg and adult tissue. The qRT-PCR results are in in line with the small-RNA-seq results (Figure 3C). To further characterize the shift from maternal-type snoRNA expression to somatic-type expression, we determined the expression of the selected maternal- and somatic snoRNAs during twelve stages of zebrafish embryogenesis. Similar to rRNAs, the predominant maternal-type snoRNAs in eggs are, during embryogenesis, gradually replaced by somatic-type snoRNAs (Figure 3B), extending the zebrafish dual ribosome system with snoRNAs.

**Figure 3.**
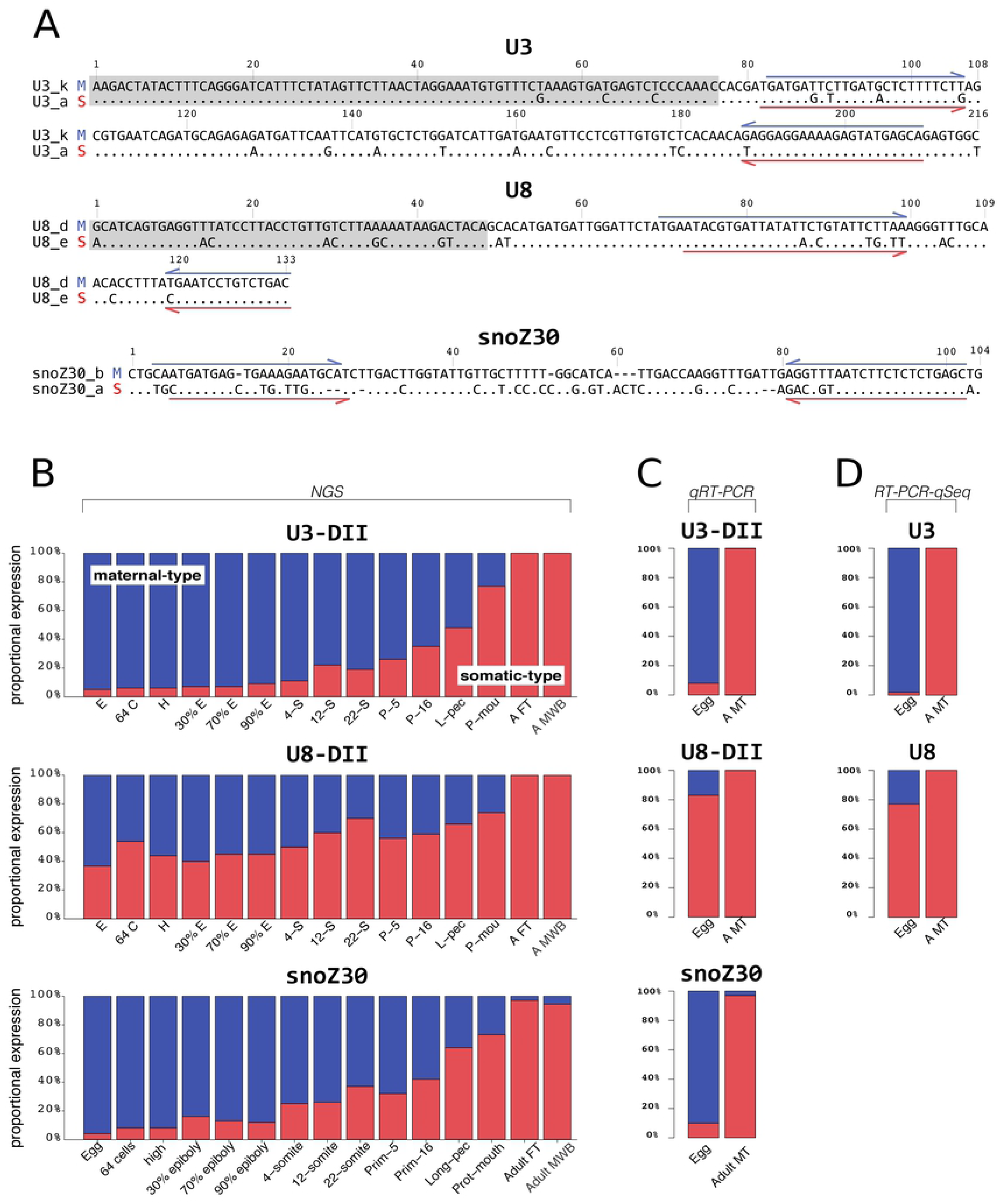
Expression of maternal- and somatic-type snoRNAs in zebrafish development. **A**: Sequence alignment of relevant maternal-type (M) and somatic-type (S) snoRNA variants; identical nucleotides are indicated as dots, while gaps as dashes (For sequence alignment of all U snoRNA cf. Supplemental Figure SF3). The qRT-PCR primers (panel C, Supplemental Table ST4) are indicated with half arrows specific for the maternal-type (blue) or specific somatic-type (red). The grey boxed sequences in the U3 and U8 snoRNAs indicate the absent regions for the observed U snoRNA-DII transcripts (Supplemental Figures SF2 and SF3). **B**: Relative expression of maternal-type (blue) and somatic-type (red) snoRNA variants indicated by comparative percentage of U3-DII, U8-DII, and snoZ30 NGS-reads, respectively. Prot-mouth: protruding-mouth; Adult FT: adult female-tail; Adult MWB: adult-male whole-body. **C**: Relative expression of maternal-type (blue) and somatic-type (red) snoRNA variants indicated by comparative percentage qRT-PCR of U3-DII, U8-DII, and snoZ30 NGS, respectively for qRT-PCR analyses on zebrafish eggs and adult male-tail (Adult MT) tissue using RT-PCR primers as indicated in panel A (Supplemental Table ST3). **D**: Relative expression of maternal-type (blue) and somatic-type (red) U3 and U8 snoRNA variants indicated by comparative percentages of RT-PCR-qSeq on zebrafish eggs and adult male-tail (A MT) tissue using RT-PCR primers and subsequent quantitative sequencing (Supplemental Table ST4).

### Structural characteristics of maternal- and somatic-types snoRNAs

A significant characteristic of the observed U3 and U8 snoRNA sequences was their length. Whereas all snoZ30 NGS-reads mapped to the whole genes (98 nt) (Supplemental Table ST1), almost all U3 NGS-reads (~130 nt.) represented just the 3’ part (Domain II (DII), Figures 1A and B), including the C/D box, of the ~215 nt U3 genes (Supplemental Figure SF3). Hence, these U3 NGS-reads are missing the U3 5’ part, including the A and A’ boxes, which guides the snoRNA in interactions with target RNAs. Consequently, the length of these U3-DII snoRNAs is quite similar to all other H/ACA box snoRNAs (Supplemental Figure SF1).

Similarly, almost all U8 NGS-reads (~86 nt) represented just the 3’ part (Domain II, Figures 1A and 4), with C/D box, of the ~131 nt U8 genes, again missing the guiding 5’ part. Whether complete U transcripts are truly absent or merely go undetected is addressed in the next section. The length of the U8-DII snoRNAs is quite similar to most C/D box snoRNAs (Table 1 and Supplemental Figures SF1 and SF3). U3-DII and U8-DII snoRNAs are reported before (13,41) and are likely a result of the removal of the mono-methylated (m^7^G) 5’ cap by the decapping complex, which is regulated by PNRC1 (42) although we cannot completely rule out that they also could originate from an alternative transcriptional start site (Figure 1A).

The U-DII snoRNAs are not merely degradation products of the complete U3 and U8 snoRNAs, as can be seen by their gradual disappearance during the embryogenesis (Figure 2B) and by the strictly defined read coverage (Supplemental Figure SF3). Thus, beside an earlier reported dominant-negative effect on ribosome biogenesis (10), we speculate that these partial U-DII snoRNAs may exert other specific functions. Since their guiding parts are missing (1,43), such function would be different from that of the traditional U snoRNA.

As a starting point for finding biological relevance of the observed maternal- and somatic-expression profiles for different snoRNAs, we compared the sequences of the maternal-type to the somatic-type variants of the U3, U8, and snoZ30 snoRNAs. There are quite some sequence differences (8%, 16%, and 36%, respectively), which may be indicative for functional diversity (Figure 3A and Supplemental Figure SF4). One obvious difference could relate to the usual target RNAs for each snoRNA.

Several potential snoRNA-rRNA interactions could be pinpointed by complementary sequences, in line with those reported in literature (1). However, many are in the 5’ guiding part of the involved snoRNAs, which is absent in our reads (Supplemental Figure SF5). Conversely, many (known) interactions are found in the ETS regions of the rRNA genes (Supplemental Figure SF5). Yet we observed that the involved maternal-type U-DII snoRNAs are present throughout embryogenesis, much like the maternal-type rRNAs. This, combined with the assumption that maternal-type rRNAs are only transcribed and processed during oogenesis and somatic-type rRNAs during embryogenesis and adulthood, it seems unlikely that maternal-type snoRNAs have a role in the processing of maternal-type rRNAs. Hence, the observed U-DII snoRNA likely have other functions, much like snoZ30 that binds to snRNA U6 (44). Given these constraints we were unable to identify any promising interactions between the many possible interactions of U-DII snoRNAs and rRNAs (Supplemental Figure SF5). It should be noted that such interactions are hard to find given the sometimes seemingly feeble reverse complement base pairing in these situations.

### Preferential expression of complete snoRNAs

The fact that virtually only U-DII snoRNA sequences were found by small-RNAseq, did not match with the current knowledge of complete U3 and U8 snoRNAs, as well as a qRT-PCR analysis which showed their abundant presence in the tested zebrafish samples (result not shown). Given that we sequenced nearly all zebrafish tissues lead to the conclusion that, with the NGS platform (Ion Proton) employed in this study, we are not able to sequence complete U3 and U8 snoRNAs. This is also the case for many 5S rRNAs plus almost all tRNAs and is likely caused by the cumulative effect of 5’/3’ modifications, strong secondary structures, and modified nucleotides.

To still determine whether the complete U snoRNAs also display preferential maternal or somatic expression, we developed a PCR-based strategy that focused on the missing 5’ parts of the U snoRNA genes. To overcome the high similarity between the U3 variants, as well as between the U8 variants, this approach starts with a RT-PCR, using as much as possible generic PCR primers (Supplemental Table ST4), after which sequencing of the PCR products reveals the distribution of variants (Supplemental Table ST4). This RT-PCR-qSeq approach revealed that the complete U3 and U8 snoRNAs display the same expression patterns as the shorter U3DII and U8DII (Figure 3D and Supplemental Table ST4).

### Possible involvement of ITS sequences in U3-DII processing

While investigating the maternal-type U3 snoRNA tandem repeats, we observed additional small RNAs that originate from the ITS regions between the snoRNA genes in the U3 genomic regions. These small RNAs either have their own promoter, or the U snoRNA genes in tandem repeats generate one transcript which is later on processed into individual mature U snoRNAs. As these small RNAs are derived from the U-ITS region, we named them snoRNA-ITS-small-RNAs (sitsRNA). For instance, for the U3 loci on chromosome 14, we detected essentially five different sitsRNAs (25 nt to 30 nt) that came from a highly conserved region of 265 nt, just 379 nt upstream of each U3 variant sequence. We noticed that there are several sitsRNAs from the various U snoRNA in tandem repeats that show complementarity with U snoRNA sequences, in particular, one 26 nt sitsRNAs from the U3 loci on chromosome 14 showed two regions (each 7 nt long) that are reverse complement to U3 snoRNA a sequence in the 5’ part and one in the D-II region (Figure 1B). This raises the intriguing possibility that this particular sitsRNAs may somehow be involved in the processing from full length U3 snoRNA to U3-DII snoRNA as the locations of interaction between the sitsRNAs and the U3 snoRNA span the cutting site (Figure 1B). Although similar sites were found in the other U3 and U8 clusters, we were unable to find another sitsRNAs that would interact like this. Still we feel there are enough indications that warrant further investigation of the possible role of U-ITS sequences in snoRNA processing.

## CONCLUSION

In this study, similar to our previously report on developmental-specific expression of rRNAs, we observed a specific subset of snoRNAs; U3, U8 and snoZ30 of which variants show distinct expression profiles during early zebrafish embryogenesis. All other snoRNAs are about eight times higher preferentially expressed in non-embryonic developmental stages, which may be a logical consequence of the fact that the rRNAs are already processed in an egg, thus only requiring snoRNAs for other tasks than rRNA maturation during early embryogenesis.

We discovered next to the complete U snoRNAs, also U3-DII and U8-DII partial snoRNAs, which miss their 5’ rRNA recognizing part and essentially consist of just the Domain-II hairpin structures. These U-DII partial snoRNAs variants also showed maternal- or somatic preferential expression, which correlated nicely with their genomic organization in tandem repeats and solitary, respectively. The complete versus partial U snoRNAs show the same preferential expression.

Though, at least part of the detected differential gene expression is likely caused by associated promotors of the involved snoRNA genes, we were unable to find any sequences in the 200 bp upstream promoter region that could discriminate the maternal-from the somatic-type snoRNA genes (results not shown).

While the function of the U-DII snoRNAs is still unclear, given that the DII part of complete U snoRNAs is known to bind to several proteins (45,46), the intact hairpin in a U-DII snoRNA hints at a role as ribonucleoprotein. In any case, since they seem to be strictly regulated, U-DII snoRNAs probably have a significant role in the zebrafish embryogenesis.

How they come about is another fascinating puzzle. We observed small RNAs, which originate from the ITS regions of U3 snoRNA loci, that show convincing complementarity with U3 sequences. This raises the possibility that they somehow may be involved in the biosynthesis of the U3-DII snoRNAs. In any case, there is a notable analogy between the relatively-small RNA-processing snoRNAs that are located in introns of genes usually involved in ribosome biogenesis and small RNAs, which are located in the ITS of snoRNAs they possibly support processing.

## MATERIAL AND METHODS

### Biological materials, RNA-isolation and small-RNA-seq

Adult zebrafish (strain ABTL) were handled in compliance with local animal welfare regulations and maintained according to standard protocols (http://zfin.org). The breeding of adult fish was approved by the local animal welfare committee (DEC) of the University of Leiden, the Netherlands. All protocols adhered to the international guidelines specified by the EU Animal Protection Directive 86/609/EEC.

For this study we used samples of two pools of unfertilized eggs (oocyte clutches) and two male-adult zebrafish tails. The harvesting of the biological materials and RNA-isolation have been described previously in (34) and (35).

### Source data

Next-generation data previously generated in our group (34) and available through the BioProject database with accession number PRJNA347637 has been used in this study with respect to i) Three pools of unfertilized eggs (oocytes); ii) one embryo at each of the 12 developmental stages: 64 cells (2 hours post-fertilization (hpf)); high stage (3.3 hpf); 30% epiboly stage (4.7 hpf); 70% epiboly stage (7 hpf); 90% epiboly stage (9 hpf); 4-somite stage (11.3 hpf); 12-somite stage (15 hpf); 22-somite stage (20 hpf); prim-5 stage (24 hpf); prim-16 (31 hpf); long-pec stage (48 hpf); protruding-mouth stage (72 hpf); and iii) one whole-body male-adult zebrafish sample.

### qRT-PCR analysis

Forward and reverse PCR primers were designed for the maternal-type snoRNA genes U3_k, U8_d and snoZ30_b, and for the somatic-type snoRNA genes U3_a, U8_e and snoZ30_a (Supplemental Table ST3). Reverse transcription was done in two independent reactions primed with the combined reverse PCR primers of either the three maternal-type variants or the three somatic-type variants. Both reactions were performed on zebrafish egg pool and whole-body adult male total RNA, in a total of four reactions. SuperScript IV Reverse Transcriptase (Thermo Fisher Scientific) was used according to the manufacturer’s instructions. Quantitative real-time PCR (qRT-PCR) was performed on 10-fold dilutions of the cDNA using a QuantStudio 3 Real-Time PCR System (Thermo Fisher Scientific).

### RT-PCR-qSeq analysis

Forward and reverse PCR primers were designed for all U3 and U8 variants, in such a way that: 1) as much as possible of the 5’-end of the full-length variants is included in the final amplicon, and 2) generic primers were selected that bind to the maternal-type, as well as the somatic-type variants (Supplemental Table ST4). cDNA was prepared as described above and used in regular PCR reactions for each of the variants independently. Amplification was performed using the Q5 High-Fidelity DNA Polymerase (New England Biolabs). The resulting amplicons were purified using the QIAquick PCR Purification Kit (Qiagen) and their size was verified on a 2200 TapeStation System (Agilent). Barcoded sequencing libraries were prepared using a modified version of the Ion Xpress Plus Fragment Library Kit (Thermo Fisher Scientific). Massive-parallel sequencing was performed on an Ion Proton System (Thermo Fisher Scientific) using an Ion PI Chip Kit v3.

### Bioinformatics analyses

#### Known snoRNA sequences

Known snoRNA sequences of *D. rerio* were downloaded from Ensemble 91 (39) and from snOPY (38) in May 2017. A union was made of these two set of sequences and sequence annotations. Characters were added to the snoRNA names to uniquely distinguish the multiple variants of the same snoRNA and a FASTA file containing only the unique snoRNA sequences was created.

#### Zebrafish snoRNA similarity tree

A hierarchical clustering was used to compare the snoRNA sequences. The dendrogram was made by using the *hclust* and *stringDist* functions from the R version 3.2.1 package ‘stats’ and ‘Biostrings’ respectively (47)

#### Mapping NGS-reads

At both 5’ and 3’ end of each snoRNA sequence, 5 Ns were added to facilitate the alignments in the NGS-read mapping. NGS-reads longer than 20 nt from all experiments, were mapped against the unique snoRNA sequences (Supplemental File SFile1) using Bowtie2 (48) with the following settings: *-np* to 0, *-- score-min* to L, −1, −0.3 for zebrafish in order to limit the maximal amount of mismatches to 5%. SAMtools v1.2 (49) was used to convert the alignment to the BAM file format and to retrieve the mapped NGS-read counts per snoRNA sequence. NGS-reads that were smaller than 50% of the length of a snoRNA sequence were discarded.

#### Analysis of NGS-read mapping results

The NGS-read counts were scaled using total raw 5.8S RNA NGS-read counts for each sample (35). The average of the three egg and the three adult-male technical replicates was taken. A cutoff of at least 100 NGS-read count for egg and adult-male combined was used to determine whether a snoRNA sequence was present and to available for the analysis of differential gene expression, which was calculated in a log2 scale of the egg over adult-male NGS-read counts.

#### Analysis of the RT-PCR-qSeq reads

A subsequence defined by the primers was selected for U3, U8 snoRNA (*Supplemental Table 4*). These subsequences were then used to search exact matching reads in the FASTQ files generated by the RT-PCR-qSeq experiments. The number of exact matches is reported as read count for each variant of U3 and U8 for all samples. The read counts belonging to the same tissues (clutch and adult male) for each variant were added up. The percentage of each variant was calculated based on total tissue reads.

#### snoRNA-rRNA interactions

BLASTn (50) was used to detect possible U3 and U8 snoRNA-rRNA base-pairing interactions starting from known interactions in human. The word-size parameter of BLASTn was set to 7 and of the resulting alignments the forward-reverse were selected with at least 10 matching base-pairs. For snoRNA-ITS interaction analysis, this was set to 7 matching base pairs.

## ACKNOWLEDGEMENTS

We are grateful for the valuable input by (anonymous) reviewers, which allowed us to improve our manuscript. We acknowledge the support of The Netherlands Organization for Scientific Research (NWO) grant number 834.12.003

## SUPPLEMENTAL FIGURES

SF1.pdf: Zebrafish snoRNA characteristics.

SF2.pdf: Tandem repeat DII snoRNA expression distribution. SF3.pdf: Maternal- and somatic-type snoRNA read coverage. SF4.pdf: Sequence alignments of snoRNA variants.

SF5.pdf: Possible snoRNA-rRNA interactions

## SUPPLEMENTAL TABLES AND FILES

SFile1.fa: Zebrafish snoRNA sequences

ST1.xlsx: Genome annotation of zebrafish snoRNAs

ST2.xlsx: Read counts from rna-seq experiment mapping to zebrafish snoRNAs ST3.xlsx: qRT-PCR primers

ST4.xlsx: RT-PCR-qSeq experiments set-up and results.

